# The human functional genome defined by genetic diversity

**DOI:** 10.1101/082362

**Authors:** Julia di Iulio, Istvan Bartha, Emily H.M. Wong, Hung-Chun Yu, Michael Hicks, Naisha Shah, Victor Lavrenko, Ewen F. Kirkness, Martin M. Fabani, Dongchan Yang, Inkyung Jung, William H. Biggs, Bing Ren, J. Craig Venter, Amalio Telenti

**Affiliations:** Human Longevity Inc., San Diego, CA 92121, USA.; KAIST, Daejeon 34141, Korea.; Ludwig Institute for Cancer Research, 9500 Gilman Drive, La Jolla, CA 92093, USA.; J. Craig Venter Institute, La Jolla, CA 92037, USA.

## Abstract

Large scale efforts to sequence whole human genomes provide extensive data on the non-coding portion of the genome. We used variation information from 11,257 human genomes to describe the spectrum of sequence conservation in the population. We established the genome-wide variability for each nucleotide in the context of the surrounding sequence in order to identify departure from expectation at the population level (context-dependent conservation). We characterized the population diversity for functional elements in the genome and identified the coordination of conserved sequences of distal and *cis* enhancers, chromatin marks, promoters, coding and intronic regions. The most context-dependent conserved regions of the genome are associated with unique functional annotations and a genomic organization that spreads up to one megabase. Importantly, these regions are enriched by over 100-fold of non-coding pathogenic variants. This analysis of human genetic diversity thus provides a detailed view of sequence conservation, functional constraint and genomic organization of the human genome. Specifically, it identifies highly conserved non-coding sequences that are not captured by analysis of interspecies conservation and are greatly enriched in disease variants.

## Main text

The estimated size of the human genome is 3.2x10^9^ base pairs. Large community and corporate efforts have identified single nucleotide variants (SNV) across the genome: 150 million SNVs in the public database dbSNP (http://www.ncbi.nlm.nih.gov/projects/SNP/snp_summary.cgi), 10 million coding variants in ExAC (http://exac.broadinstitute.org) and 170 million SNVs in 10,545 deeply-sequenced whole genomes ^1^ (http://hli-opensearch.com). The union of these resources is 242 million unique SNVs - representing the current reporting of single nucleotide variable sites across the whole human genome. This suggests that 1 out of every 13 nucleotides in the genome has an observed variant in the population.

There is a good understanding of protein-coding variants – owing to historical studies of Mendelian disorders, the predictable consequences of amino acid change, and the recent availability of exome sequencing data ^2^. However, the protein-coding regions represent less than 2% of the total genome, and relatively little is known about the functional consequence of variation in the remaining 98% of the genome. The non-protein-coding sequences of the genome (thereafter in this text described as “non-coding”) have been annotated through the ENCODE project that relies on identification of biochemically active elements in the human genome, with particular attention paid to regulatory elements that control gene activity (https://www.encodeproject.org). Regulatory control can also be influenced by higher-order chromatin structure, such as long-range chromatin loops ^3^, that regulates the accessibility and proximity of genes and regulatory elements. Supporting a role for non-coding variants in human disease and phenotypic traits, most of the over 16,000 common variants identified through genome-wide association studies (GWAS) at p<5x10E-7 are in non-coding regions of the genome (http://www.ebi.ac.uk/gwas). GWAS variants are increasingly recognized as acting through changes in the regulatory circuitry ^4,5^. Consistent with this hypothesis, a subset of variants is also specifically characterized as expression quantitative trait loci (eQTL) through defined *cis* or *trans* association with expression levels of gene transcripts ^6^. Despite recent progress in the study of non-coding variants, it remains a significant challenge to characterize the non-coding variants in the human genome, which grow by over 8,000 with each additional genome sequenced ^1^.

To better characterize the population variation in the non-coding regions we performed a comprehensive analysis of 11,257 whole genome sequences. We applied a metaprofiling approach ^1^ that exploits the multiplicative contribution of elements in thousands of genomes. Metaprofiles integrate and score sequence variation and frequency across genomic landmarks sharing the same sequence, structure or function. In the present work, we extend this approach by using massive alignments of k-mers to determine the probabilities of variation of each nucleotide genome-wide in the context of the surrounding nucleotides. Specifically, we exploit heptamers (7-mers) for the analysis; the heptanucleotide context was shown recently to explain >81% of variability in substitution probabilities ^7^.

The 16,384 unique heptamers present in the human genome vary greatly in abundance, ranging between 1,927 and 6,314,598 counts per genome (Suppl. Fig. S1). Heptamers are not evenly distributed across the genome and some show clear association with genomic elements (**Fig. 1A**). Each heptamer is characterized by unique rates of variation. To capture this property, we computed the rate and frequency of variation at the 4th nucleotide of each different heptamer (see Methods). The metric varies > 100-fold across heptamers (between 0.0015 and 0.157; Suppl. Fig. S1). It defines the expectation of variation for each nucleotide in the genome.

**Figure 1.**
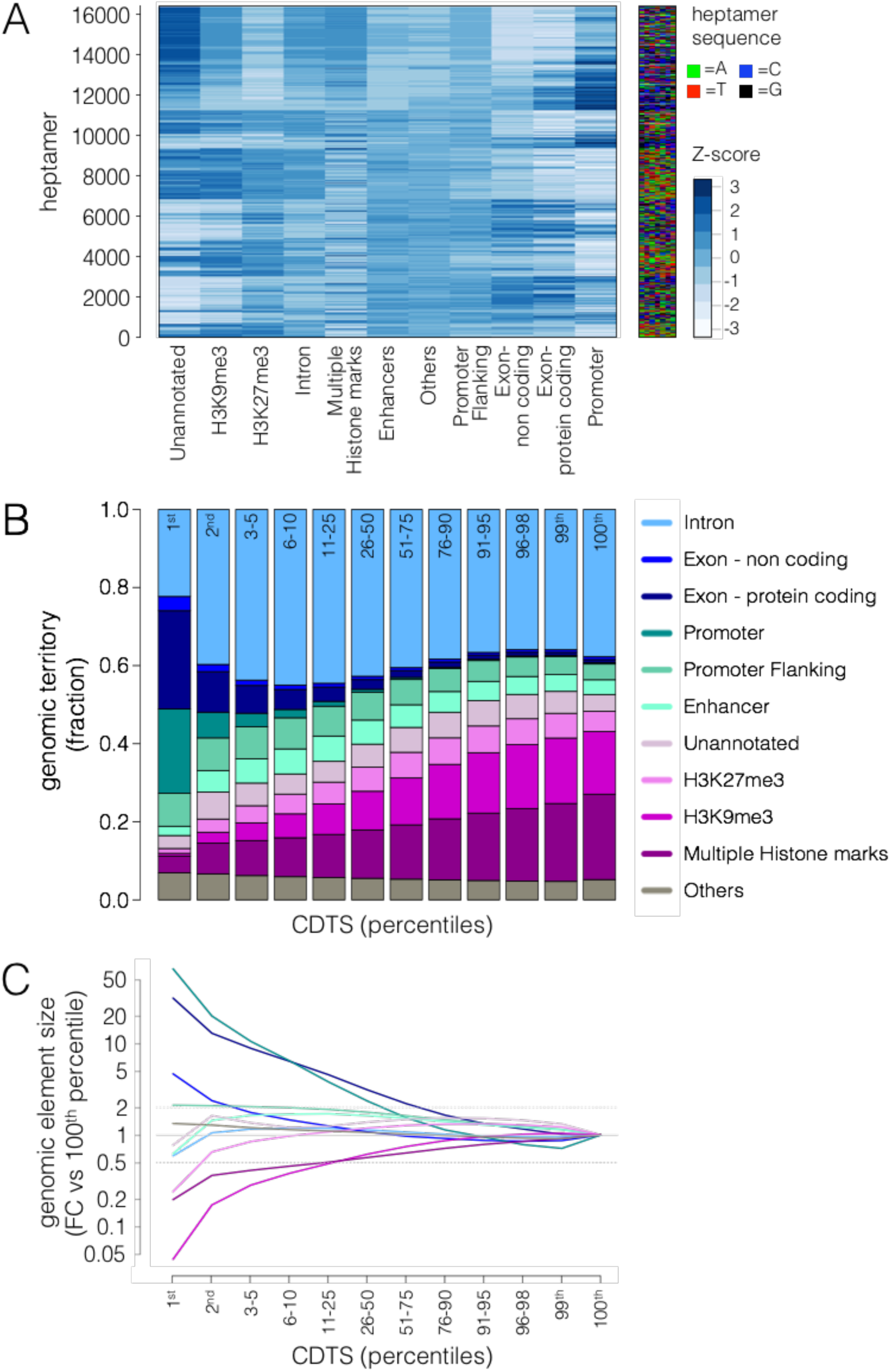
K-mer structure of the genome and composition of the conserved human genome. **A.** The blue shades heatmap represents the relative composition of k-mers for the different genomic elements. Each row corresponds to a heptamer, with the corresponding nucleotide sequence displayed on the heatmap to the right. The relative abundance of a heptamer should be compared horizontally across the genomic element with the shades of blue reflecting the z-score. Before standardization, the counts of heptamers have been normalized per genomic element to take into account the different territory sizes of the element families. The order of the rows was obtained by hierarchical clustering. **B**. The barplot displays the cumulative territory fraction covered by each element family in the different percentile slices (indicated at the top of the bars). Here, and in other figures, we purposefully emphasize the patterns a the lowest 1^st^, 2^nd^, and 3-5^th^ percentiles. The percentiles are based on the rank of CDTS values. “Others” refers to ENCODE element families that did not cover a substantial part of the genome individually (such as transcription factor binding sites, see Methods). The elements appear in the same order as in the legend. **C**. The enrichment and depletion of each percentile slice compared to the 100^th^ percentile. The fold change is normalized by the size of the slice. Element families are colored as in panel B. CDTS, context-dependent tolerance score. FC, fold change. Vs, versus.

A given heptamer or region may have rates of observed variation that are higher or lower than rates estimated genome-wide. We defined the context-dependent tolerance score (CDTS) as the absolute difference of the observed variation from expected variation. Thereafter, we divided the genome into equal size regions using a sliding window of 50 base pairs (bp) to study the context-dependent conservation without consideration of existing annotation. Based on CDTS we rank every region in the genome from the most context-dependent conserved (1^st^ percentile) to the least context-dependent conserved (100^th^ percentile) (**Fig. 1B** and Suppl. Fig. S2A). We identified patterns of enrichment and depletion for specific genomic elements as well as chromosomes across the spectrum of CDTS values (**Fig 1C**, Suppl. Fig. S2B; See Methods for the categorization of the genomic elements). As expected, protein coding exons were strongly enriched (31-fold) in the first percentile of CDTS. Specifically, 11,901 protein coding genes had at least 1 exon in the first percentile; only 1,816 genes had no single exon in the first 10 percentiles. The context-dependent correction also identified a striking enrichment for promoters (66-fold) in the most conserved regions of the genome. Despite no clear enrichment or depletion pattern of enhancers throughout the CDTS spectrum, super-enhancers were enriched at lower CDTS, in a magnitude proportional to the number of cell types they were present in. (Suppl. Fig. S2C). In contrast, marked depletion was observed for H3K9me3 histone marks (23-fold). However, it is important to underscore that all families of genomic elements, as well as unannotated genomic sequences are found in the most context–dependent conserved regions of the genome as measured by CDTS (**Fig. 1B**, Suppl. Fig. S2D). To compare these findings in the larger context of interspecies conservation, we assessed the extent of overlap of conserved regions assessed with CDTS (ie., context-dependent conservation in the current human population) and Genomic Evolutionary Rate Profiling (GERP) across 34 mammalian species (ie., interspecies conservation). From the 1^st^ to 10^th^ percentile levels, the overlap between both scores is limited and heavily enriched for protein coding regions (Suppl. Fig S3). Taken together, these results suggest that the most constrained non-coding regions in human populations are primarily revealed by CDTS.

A large proportion of the thus defined constrained human non-coding genome is associated with regulatory elements such as promoters, enhancers, transcription factor binding sites and regions associated with active chromatin marks (**Fig. 1B**). We hypothesized that the most constrained regulatory regions serve to regulate the most functionally important genes. To test the hypothesis, we used the notion of gene essentiality to define “functional importance”. Essential genes are characterized by limited tolerance to truncation, fewer paralogs, and being part of larger protein complexes ^1,2,8^. Editing of those genes with CRISPR-Cas9 compromise cell viability, and knockouts of these genes in the mouse model are associated with increased mortality ^8^. As expected, exons in essential genes were enriched in the conserved regions of the genome as defined by CDTS (**Fig. 2A**). Thereafter, we assigned the essentiality score of the gene to the corresponding upstream promoter. This analysis confirmed that promoters in the constrained part of the genome associate with essential genes (**Fig. 2A**). We then observed that *cis* enhancer regions also shared sequence constraint with genes (within 15 kb) that were putatively regulated by those elements (**Fig. 2A**). Next, we searched for evidence that functional constraints could be shared over greater distances. Topological associated domains were defined using information from promoter capture Hi-C (pcHi-C), Hi-C and 3D genome structure data ^9–11^. We observed that the regions brought together through these long-distance interactions shared similar levels of conservation as reflected by the CDTS values. This coordination was maintained at distances as long as one megabase (Mb) (**Fig. 2B**, Suppl. Fig S4). In addition, with the newly developed pcHi-C technique^11^, enabling to associate distant regulatory regions with a particular gene, we observed a correlation between conservation of the distal enhancer, and the essentiality of the target gene (**Fig. 2C**; see Methods for pcHi-C technical description). Finally, we assessed other *cis* non-coding elements (eg., chromatin histone marks, transcription factor binding sites), unannotated and intronic regions, and consistently identified a pattern of correlation between CDTS of non-coding or regulatory regions with gene essentiality (**Fig. 2A**). Strikingly, even genomic elements that were depleted in the most conserved part of the genome (e.g. H3K9me3 and H3K27me3) are associated with essential genes when present in the lower CDTS percentiles (**Fig. 2A**). More generally, regions of low CDTS appear clustered in the genome (Suppl. Fig S5). Overall, the data support the concept of constrained and coordinated regulatory and coding units in the genome over large genome distances.

**Figure 2.**
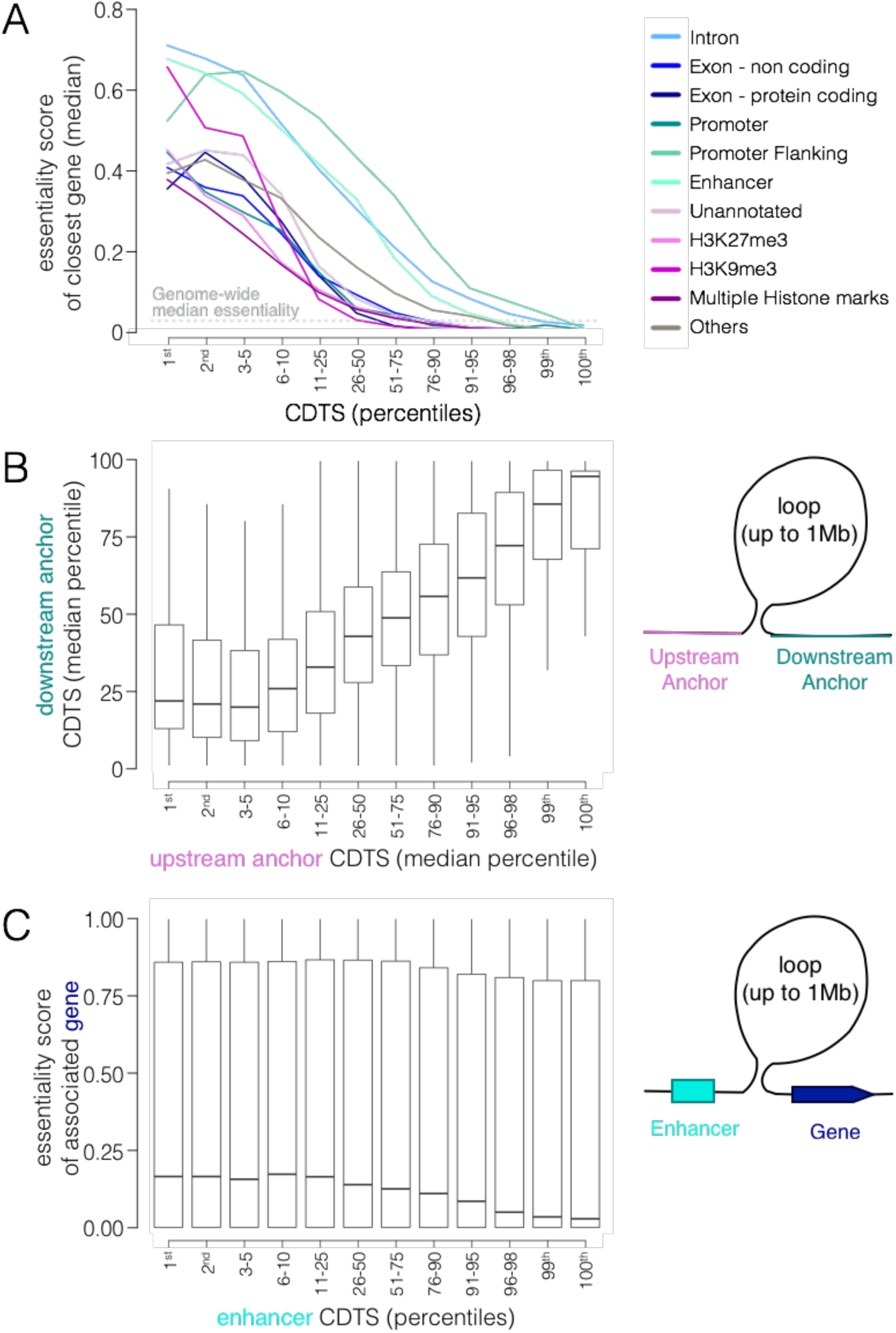
Shared conservation of genes and *cis* or distal regulatory elements. **A.** Coordination of *cis*-elements. Each genomic bin within 15 kb of a gene (*cis*) is attributed the essentiality score of the closest gene. The median essentiality score of the closest genes is depicted on the Y-axis for each genomic element family throughout the CDTS spectrum (X-axis). The grey horizontal dashed line represents the median gene essentiality score genome-wide (0.028). **B.** Distal coordination of anchor regions. A chromatin loop is depicted in the right panel. The median CDTS is extracted for each anchor region and binned in percentile slices. The X- and Y-axes indicate the median CDTS values for the upstream and downstream anchor regions, respectively. The anchor regions surrounding a loop share CDTS values. The whiskers extend from the 10^th^ to the 90^th^ percentiles of the data. The box spans the interquartile range. Outliers are not displayed. **C.** CDTS-essentiality coordination of promoter distal regions and their putative target gene pairs obtained from promoter-capture Hi-C results (n=483,517) ^11^.The boxplots depict the essentiality of the putative target genes (y-axis) along the distal region CDTS (x-axis). The whiskers extend from the 10^th^ to the 90^th^ percentiles of the data. The box spans the interquartile range. Outliers are not displayed. CDTS, context-dependent tolerance score.

The description of the conserved regulatory units raises the issue of its relevance to human disease. We assessed whether CDTS ranking was a good proxy to score functionality and the consequences of mutations. For this purpose, we investigated the distribution of annotated pathogenic variants across the genome. The pattern of enrichment was marked for pathogenic variants in the 1^st^ versus the 100^th^ percentile for both protein-coding (77-fold) and, more importantly, for non-coding (80-fold) pathogenic variants (**Fig. 3A**). Of note, the enrichment of non-coding pathogenic variants is even more striking after accounting for the size of the non-coding territory covered in each percentile slice and reaches > 110-fold enrichment (Suppl. Fig. S6). To confirm these findings, we further investigated 550 manually curated non-coding variants associated with Mendelian disorders^12–15^. We confirmed that Mendelian non-coding variants are highly enriched in the regions with the lowest CDTS values (**Fig. 3B**). The nature and disease association of variants and their CDTS values are summarized in Table S1.

**Figure 3.**
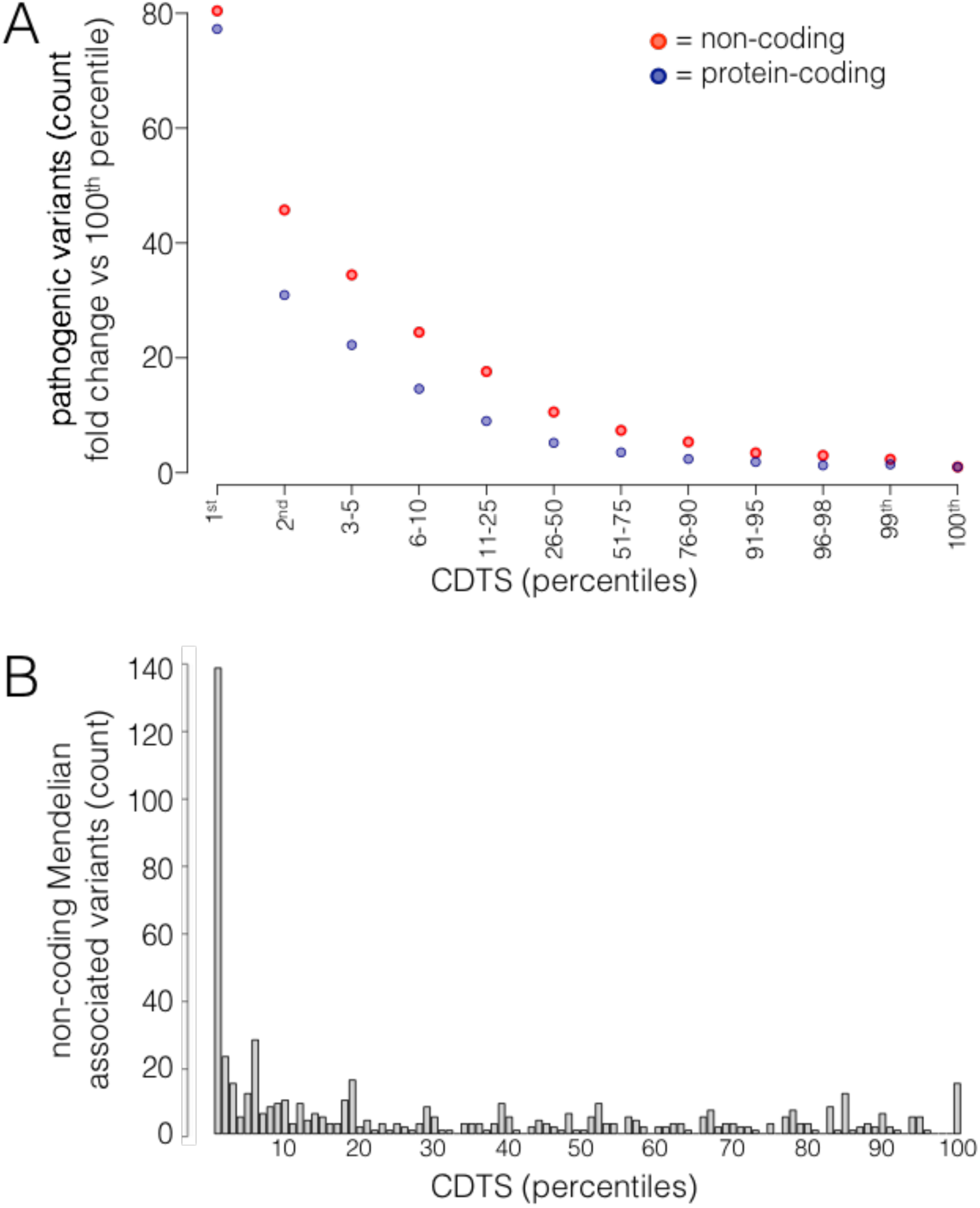
Distribution of pathogenic variants across the genome. **A.** The distribution of pathogenic variants across the different percentile slices identifies a strong enrichment at lower CDTS percentiles. The relative enrichment is calculated with regards to the 100^th^ percentile. Protein-coding pathogenic variants are shown in dark blue; non-coding pathogenic variants in red. The total number of pathogenic variants are N=120,759 protein coding and N=14,092 non-coding variants. Exonic non-coding (e.g. lincRNA) are not displayed as only a very limited number of pathogenic variants were annotated (N=557). **B.** Non-coding pathogenic variants associated with Mendelian traits. The total number of Mendelian associated non-coding pathogenic variants is N=550. Pathogenic variants are enriched at the lowest percentiles. CDTS, context-dependent tolerance score. Vs, versus.

We explored how CDTS compared to other functional predictive scores used to prioritize variants, such as CADD and Eigen ^16,17^. We focused on the performance of these metrics on the non-coding genome. The combination of the three metrics provides the best detection, while the three metrics used alone provide similar ranges of detection (**Fig. 4A**). CDTS is the functional predictive score that has the highest fraction of specific variant detection at any percentile threshold (**Fig. 4B**, barplot) providing high complementarity to the other metrics, while Eigen and CADD capture more redundant information (**Fig. 4B**, Venn diagrams). In addition, CDTS is the functional predictive score that detects the highest number of pathogenic variants, as the scores are computed for the whole genome, including sex chromosomes, and can be used for both SNVs and indels (**Fig. 4B**, Venn diagrams). Overall, CDTS requires no *prior* knowledge such as annotation or training sets, and captures a very specific set of pathogenic variants that are not detected by other metrics. Thus, CDTS complements other functional predictive scores in the analysis of the non-coding genome.

**Figure 4.**
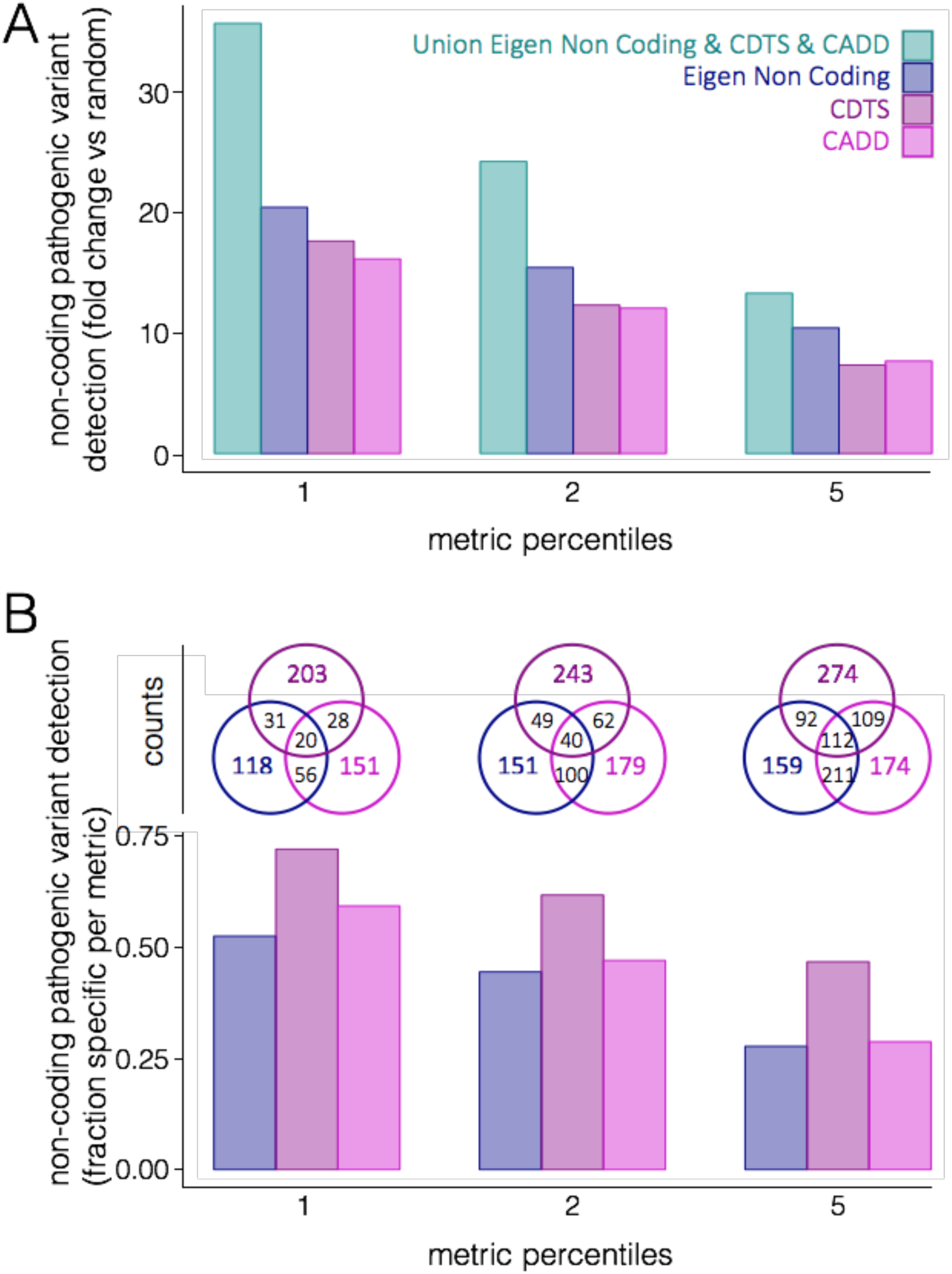
Complementarity of scores for non-coding variants. **A.** The enrichment of pathogenic variant detection, as compared to random, is displayed at different percentile thresholds for Eigen non coding, CDTS, CADD as well as for the union of the three metrics. **B.** The barplot displays, at different percentile thresholds, the fraction of pathogenic variants identified exclusively by only one of the metrics. The Venn diagram displayed on top of each percentile threshold shows the overlap of pathogenic variant detection for Eigen non coding, CDTS and CADD. CDTS, context-dependent tolerance score. CADD, combined annotation dependent depletion. Vs, versus.

In summary, we assessed conservation of the human genome solely based on human variation. The analysis first established the expectation of variation based on sequence context. The approach identifies regions of the genome, that while having various levels of absolute variation, are nonetheless under selective constraint. Its clinical relevance is manifested by the enrichment of known pathogenic variants in the most constrained part of the genome. A practical implementation of this observation is the targeting of sequencing efforts beyond the exome. Many exons could possibly be eliminated from targeted analysis while including an equivalent amount of sequence that represents the most constrained regions of the non-coding genome. The second important observation is the complementarity of human conservation metrics with other analyses of the non-coding genome. Kellis et al. ^18^ reviewed the contribution of biochemical, evolutionary (interspecies conservation) and genetic approaches for defining the functional genome. They concluded that each approach provided complementary information and that the combination of approaches was most informative. Our data indicates that CDTS based on human diversity, serves as a fourth approach for the characterization of the non-coding human genome. The last and most biologically important observation is the organization of functional units to share a conservation profile. The data indicates that an essential gene will use proximal and distant regulatory elements that are co-conserved. Use of this information supports the identification of *cis* or distal rare variants that regulate the expression of medically important genes.

## Methods

### Genomes

The analysis used deep sequence genome data of 11,257 individuals. Analysis was limited to the high confidence region of the genome as defined in Telenti et al. (1) - a region covering approximately 84% of the genome and closely overlapping with the high confidence region as described in the most recent release of Genome in a Bottle (GiaB v3.2, ftp://ftp-trace.ncbi.nlm.nih.gov/giab/ftp/release/NA12878_HG001/NISTv3.2.2/).

### Metaprofiles

Metaprofiles consist of the massive alignment of elements of the same nature in the genome ^1^. These genomic elements can be chosen based on their structure (e.g., exonic, intronic, intergenic, etc.), function (e.g., transcription factor binding sites, protein domains, etc.) or sequence composition (*k*-mers). Genetic diversity is assessed at each nucleotide position of the alignment of genomic elements, by monitoring both the occurrence of variation in the population (reported as a binary – presence or absence) and the allelic frequency. More specifically, 3 metrics are computed at each position: (i) the percent of elements with SNVs (count score; Suppl. Fig S1B), (ii) the percent of SNVs with an allelic frequency higher than 0.001 (frequency score; Suppl. Fig S1C), and (iii) the product of both scores (tolerance score; Suppl. Fig S1D). Each score is calculated using between 10^6^ and 10^10^ values, a value provided by the number of elements present in the genome and aligned multiplied by the number of genomes sequenced; therefore, the metaprofile strategy massively increases the power to compute variation rate at nucleotide resolution with high precision. *A priori* knowledge of genomic landmarks is required for constructing metaprofiles based on similarity in structure or function. In order to remove potential biases through the use of this *a priori* knowledge, we developed a strategy to construct metaprofiles based on all possible heptameric sequences found in the genome (4^7^=16,384) and scored the middle nucleotide for each of these sequences as described above. As every nucleotide in the genome is part of an heptamer, every single position can be attributed to the corresponding genome-wide computed scores.

### Expected versus observed

The variation rates computed through heptamer metaprofiles reflect the chemical propensity of a nucleotide to vary depending on its surrounding context and can be interpreted as an expectation of variation. We rationalized that functional regions would vary significantly less than they would be expected to, as assessed genome-wide through the heptamer tolerance score. To evaluate the departure from expectation, we compared the observed and expected tolerance score obtained in defined genomic regions.

The observed regional tolerance score is the number of SNVs present at an allelic frequency higher than 0.001 in the studied population in a defined region. The expected regional tolerance score is the sum of the heptamer tolerance scores in the same region.

The difference between the observed and expected scores is further referred to as *context-dependent tolerance score* (CDTS). The regions are then ranked based on their CDTS. The regions with the lowest rank (1^st^ percentile) are the regions with the lowest context-dependent tolerance to variability and the regions with the highest rank (100^th^ percentile) are the regions with the highest context-dependent tolerance to variability.

### Region definition and annotation

To avoid any use of *a priori* knowledge and any biases due to the differing size of the regions (e.g., more power to detect difference between observation and expectation in longer elements), the genome was chopped irrespective of genomic annotations into sliding windows of the same size. The window size was 1050 bp sliding every 50 bp and the calculated CDTS across the 1050 bp window was attributed to the middle 50 bp bin. Only regions with at least 90% of the nucleotides in the 1050 bp window present in high confidence regions were used. To evaluate the element distribution across those size defined windows, we built a new annotation model by combining sources of annotation from GenCode (v.23) and ENCODE (annotated features and multicell regulatory elements, Ensembl v84 Regulatory Build). In order to avoid conflicting and overlapping annotations from the two different sources and thereby use the score of the same region multiple times, we prioritized element annotation as follows, such that only the highest order element would be used: exonic, then multicell, then intronic and then annotated features. We assessed the element composition of the different percentiles, using the above mentioned combined GenCode/ENCODE annotation, by computing the number of nucleotides of an element in each percentile. The following categories were used: “Exon - protein coding”, referring to nucleotides in exonic regions contained in protein coding genes (including UTR) as annotated in GenCode; “Exon - non coding”, referring to nucleotides in exonic regions contained in non-coding RNAs (e.g., snRNA, snoRNA, lincRNA, etc.) as annotated in GenCode; “Intron”, referring to nucleotides in intronic regions contained in either protein coding or non-coding genes as annotated in GenCode; “Promoter”, “Promoter Flanking” and “Enhancer”, referring to the nucleotides contained in the respective elements as annotated in ENCODE multicell regulatory elements; “H3K9me3” and “H3K27me3”, referring to the nucleotides overlapping with (and only) the respective elements as annotated in ENCODE annotated features; “Multiple Histone marks”, referring to the nucleotides overlapping with a combination of histone marks, as annotated in ENCODE annotated features; “Others”, referring to the remaining nucleotides with ENCODE multicell regulatory element or annotated features that did not cover a substantial part of the genome individually, which notably encompasses transcription factor binding sites as well as other regulatory element combinations (e.g., nucleotides annotated as both Promoter and Enhancer); and “Unannotated”, referring to nucleotides in regions that had no annotation in either GenCode or ENCODE. Super-enhancer annotation (in Suppl. Fig. S2C) was obtained from dbSUPER (http://bioinfo.au.tsinghua.edu.cn/dbsuper/index.php).

### Essentiality and CDTS coordination

We used gene essentiality (pLI score from ExAC ^2^) as an orthogonal proxy for functionality to assess whether genomic bins, annotated with the same genomic element, have different biological importance depending on their CDTS ranking. Each genomic bin present within 15 kb of a gene is attributed the essentiality score of its closest or overlapping gene, with the exception of genomic bins annotated as “Promoters”, that have the mandatory constraint of being upstream of the closest gene. The median essentiality score is then assessed per genomic element annotation and per percentile slice. To assess distal CDTS coordination, we used two external datasets. To test the possible coordination of anchor regions, we used a Hi-C dataset aggregating the results of multiple cell types^10^. The median CDTS percentile is computed for every anchor region. To test the possible coordination of distal gene-enhancer pair, we used a pcHi-C dataset performed in lymphoblast cell lines. ^11^ in order to identify promoter-centered long range interactions. Briefly, pcHi-C library was constructed by performing a target enrichment protocol (enriching target promoter-centered proximity ligation fragments from Hi-C library using capture RNA probes). Unmapped, non-uniquely mapped, PCR duplicates, trans-chromosomal read pairs, putative self-ligated products (<15kb read pairs), and off-target reads were removed. We further eliminated experimental biases by using “capture” scores, a probability of the region being captured. We only considered promoter-centered long-range interactions within the distance of 2Mb from the transcription start site. Significant pcHi-C chromatin interactions were identified in terms of p-value 0.002 cutoff of Weibull distribution after removing distance dependent background signals. After filtering, the CDTS of each remaining so-called distal “interacting” genomic bin is compared to the essentiality score of the interacting gene.

### Interspecies conservation

We used Genomic Evolutionary Rate Profiling (GERP++) ^19^ to capture the interspecies conservation. GERP++ provides conservation scores through the quantification of position specific constraint in multiple species alignments. We calculated and attributed the mean GERP scores to the same set of 50 bp bins as mentioned in the section “*Region definition and annotation*”. Bins were ranked based on the GERP score from the most (percentile 1) to the least conserved (percentile 100). Bins without GERP score, due to insufficient multiple species alignments in the region, were not considered in the ranking process.

### Pathogenic variants

We assessed the distribution of known annotated pathogenic variants, defined as either HGMD high DM ^20^ (Version: HGMD_2016_R1) or ClinVar variants consistently annotated as pathogenic or likely pathogenic and with at least 1 entry with star 1 or more^21,22^ (Version: ClinVarFullRelease_2016-07.xml.gz) for a total N=135,965, by counting the number of variants present in each percentile of the genome. For variants in indel regions, the left most coordinate was used to establish in which genomic bin they fell. Pathogenic variants with conflicting annotations were removed, defined here as variants having a high DM in HGMD and a consistent annotation of benign or likely benign with at least 1 entry being star 1 or more in ClinVar. The non-coding variants associated with Mendelian traits were extracted from ClinVar (copy number variants were excluded from analysis) and manually curated and additional variants were collected by literature review ^12–15^. Variants falling in untranslated regions (UTR) were considered as non-coding for the purpose of the Mendelian pathogenic associated variants. A filter of >5 bp from any splice acceptor or splice donor site was applied, as splice sites variants have a high likelihood of being pathogenic and would not be a good control to test our model. Of note, intronic bins that have the lowest CDTS are more likely to be in the extended surrounding of splice sites, raising the possibility that it might be equally important to keep the surrounding sequence conserved (Suppl. Fig. S7).

### Functional predictive scores

The CDTS metric was compared to the most widely used metrics for variant prioritization: CADD ^16^ and Eigen ^17^. A “control” set of variants relative to the previously defined pathogenic variants was created using variants from dbSNP ^23^ (June 2015 release). A control variant was defined as having the “COMMON” and “G5A” tag (>5% minor allele frequency in each population and all populations overall) and, similar to the tested pathogenic variant set, not be present in an exonic region and appear more than 5 bp from any splice site. The remaining working set of non-coding pathogenic and control variants were ranked according to their CDTS, CADD or Eigen NonCoding scores and the ranking was normalized from 0 to 100 (for CADD and Eigen, the PHRED scores were converted into probabilities before this step, so that for all metrics the lower the ranking the more likely pathogenic a variant would be). To compare the different metrics, the precision (TP/(TP+FP)) was computed at each step of the new ranking. TP are the true positives, in this case the number of pathogenic variants with a ranking ≤*threshold*, and FP are the false positives, in this case the number of control variants with rank ≤*threshold*; where *threshold* can be any step in the new ranking (from 0 to 100). For the union of the 3 metrics, all variants (pathogenic and control) that had a score in at least 1 metric were used. TP represented the number of pathogenic variants with a ranking ≤*threshold* in 1 or more of the metrics; and FP, represents the remaining number of variants to reach the number of variants present at *threshold*: (((∑pathogenic+∑control)/100) * *threshold*) - TP. The precision was further normalized by the general prevalence of pathogenic variant in the set studied (∑pathogenic/(∑pathogenic+∑control)). This step was done in order to account for the fact that not all variants were scored by the other metrics (e.g., no scores on chromosome X for Eigen, conversion conflicts from hg19 to hg38, not all indel have a CADD score, etc.). The prevalence normalized precision provides the enrichment of a metric pathogenic variant detection compared to random.

### Data access

The genome-wide CDTS values are available through hli-opensearch.com. Use of the query terms CDTS1 and CDTS10 will return all variants with CDTS scores within the 1^st^ and the 10^th^ percentile, respectively.

## Supplementary materials

**Supplementary Figure 1.**
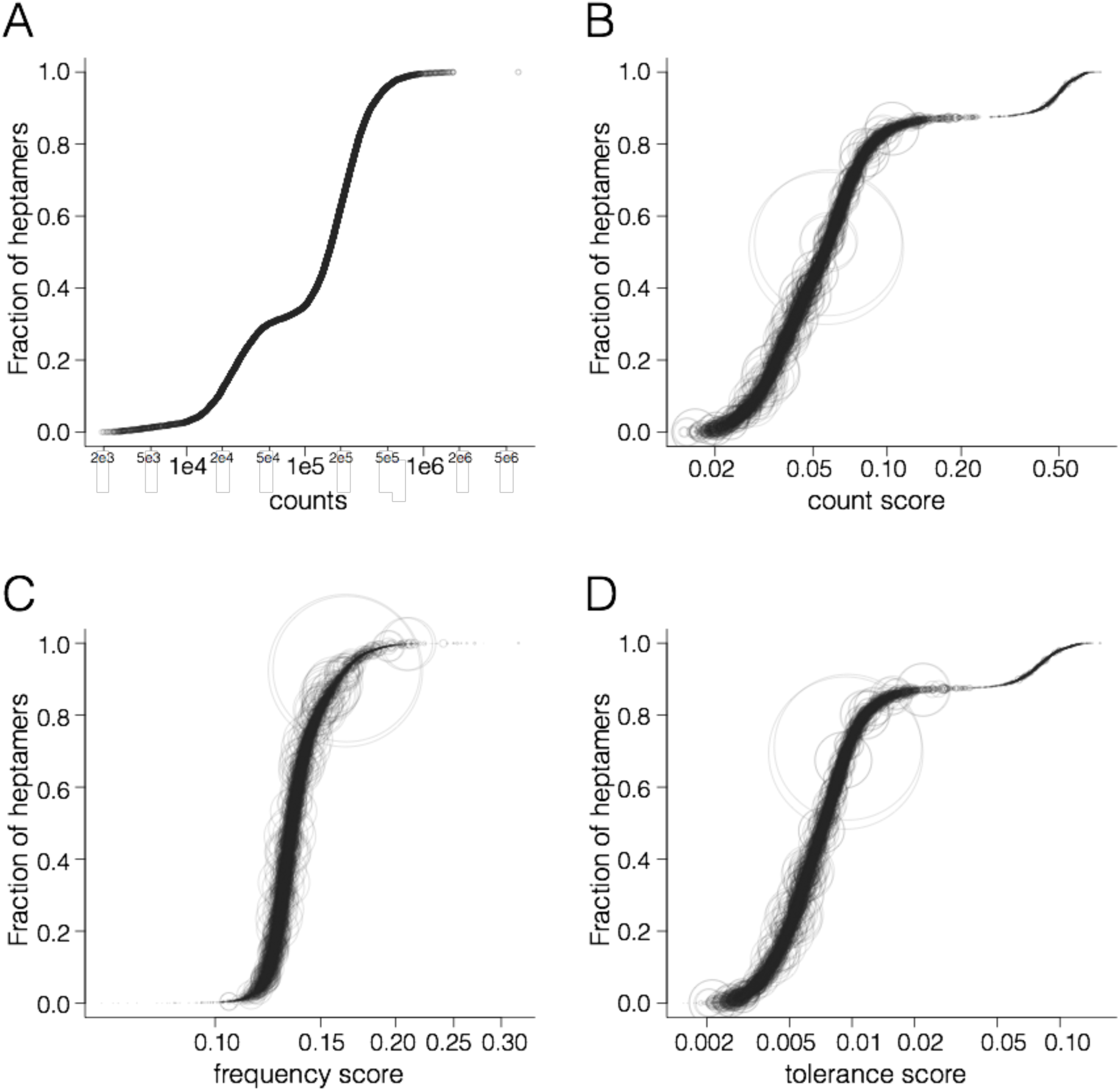
Heptamer metrics in the human genome. **A.** Cumulative distribution function of the total number of occurrence of each heptamer in the genome. Each dot represents an heptamer. Heptamers are ranked by the number of occurrence in the genome. **B.** Cumulative distribution function of the count scores. The count score represents the fraction of the middle nucleotide in a heptameric sequence that varies. Every circle represents an heptameric sequence. The heptamers are ranked by their count score. The size of the circles is proportional to the number of occurrences of the heptamer in the genome (plotted in panel A.) **C.** Cumulative distribution function of the frequency scores. The frequency score represents the fraction of SNV at the middle nucleotide in a heptamer that varies with an allelic frequency > 0.001. Every circle represents a heptameric sequence. The heptamers are ranked by their frequency score. The size of the circles is proportional to the number of occurrences of the heptamer in the genome (plotted in panel A.). **D**. Cumulative distribution function of the tolerance scores. The tolerance score represents the probability of the middle nucleotide in a heptamer to vary with an af > 0.001. Every circle represents an heptameric sequence. The heptamers are ranked by their tolerance score. The size of the circles is proportional to the number of occurrences of the heptamer in the genome (plotted in panel A).

**Supplementary Figure S2.**
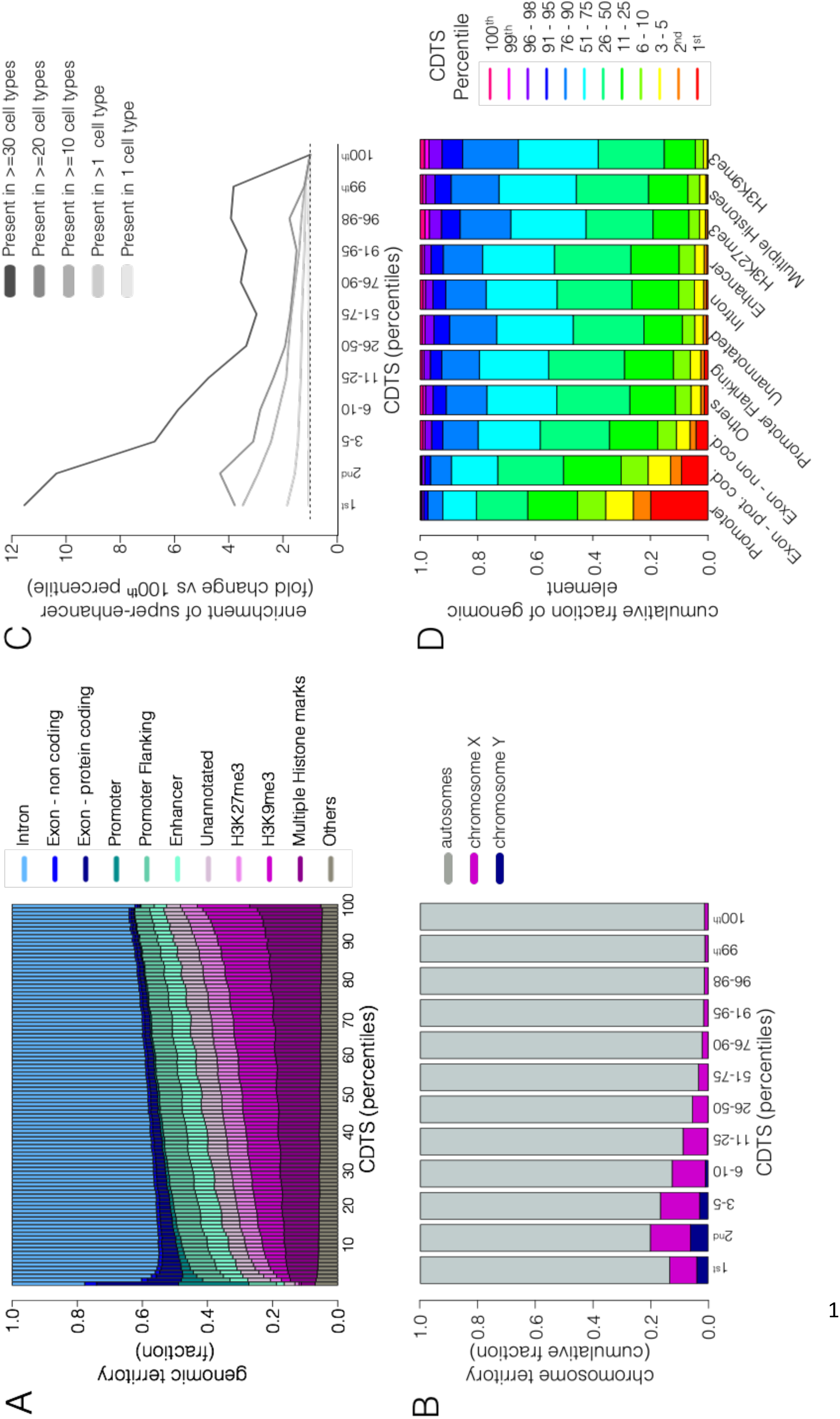
Distribution of genomic element and chromosomes within CDTS spectrum. **A.** The barplot displays the cumulative territory fraction covered by each element family in the different percentile (1 to 100). “Others” refers to ENCODE element families that did not cover a substantial part of the genome individually (such as transcription factor binding sites, see Methods). The elements appear in the same order as in the legend. **B.** The barplot displays the cumulative territory fraction covered by autosomes and sex chromosomes. Sex chromosome are enriched in the regions with lower CDTS. **C.** Size normalized distribution of super-enhancer annotation. The relative enrichment of the fraction of enhancer bins overlapping with super-enhancer annotation is calculated with regards to the 100^th^ percentile. Super-enhancers were sub-categorized depending on the number of cell types they were annotated in, represented by the multiple shades of grey lines. **D.** The barplot displays the distribution of the total amount of nucleotides within the percentile slices for each element family. The boxes within a bar indicates the fraction of element in a given percentile slice (e.g., 20% of the promoters are within the 1^st^ percentile). The element families are ordered on the X-axis by the fraction of element within the 1^st^ percentile slice. The coloring of the boxes is in the same order as the legend. Prot., protein. Cod., coding. CDTS, context-dependent tolerance score.

**Supplementary Figure S3.**
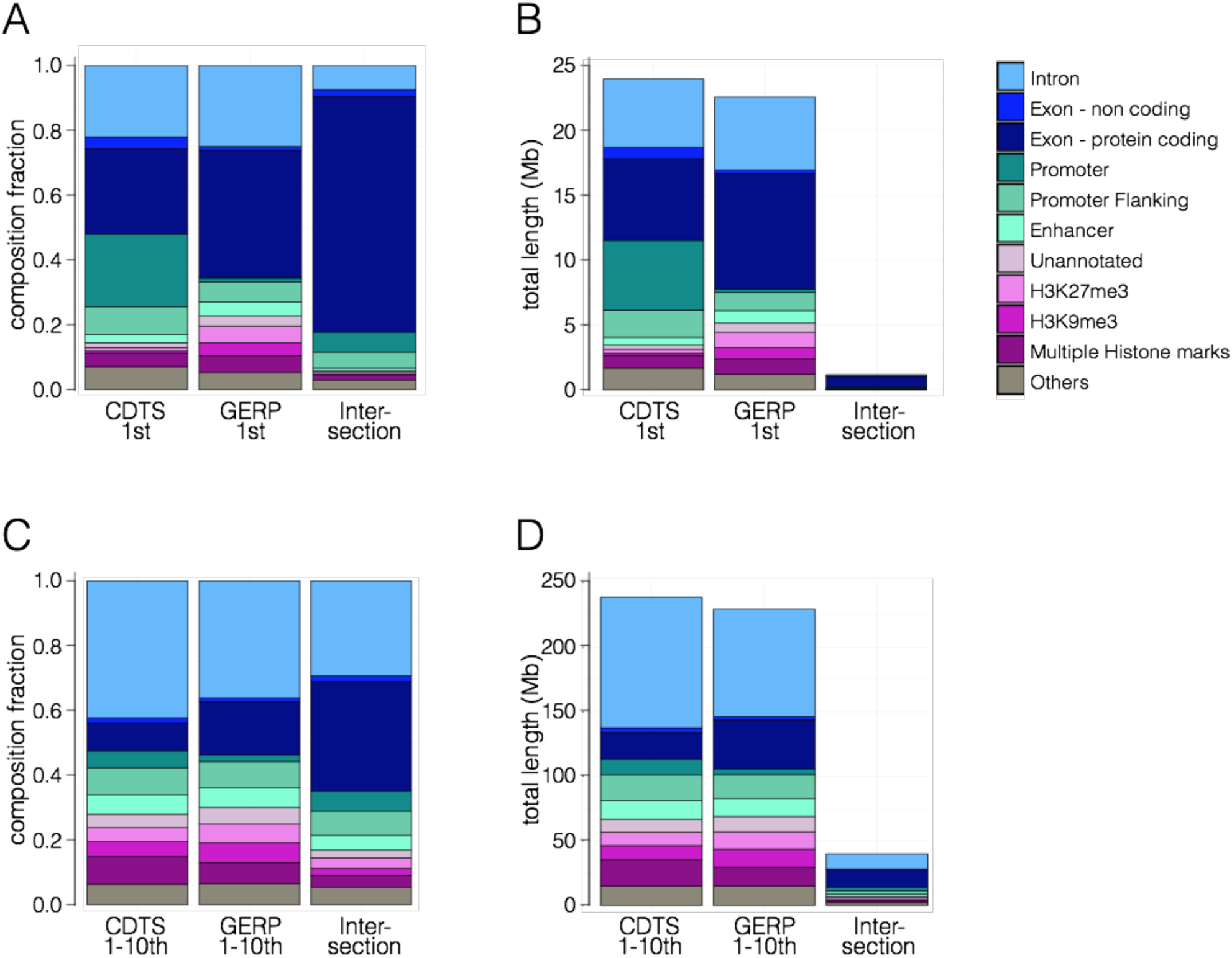
Comparison of conserved regions assessed with CDTS and GERP. **A**. Element family composition in the first percentile regions of CDTS (the bar labelled as “CTDS 1^st^”), GERP (“GERP 1^st^”) and the overlap region of CDTS and GERP (“Intersection”). Boxes in the bar correspond to different element families. The coloring of the boxes is in the same order as the legend. **B**. Length of the first percentile regions of CDTS, GERP and the overlap region of CDTS and GERP. Bins without GERP score, due to insufficient multiple species alignments in the region, were not considered in the ranking process. This explains the total length difference between the first percentile regions of CDTS and GERP. **C.** Element family composition in the first 10 percentile regions of CDTS (the bar labelled as “CTDS 1-10^th^”), GERP (“GERP 1-10^th^”) and the overlap region (“Intersection”). **D.** Length of the first 10 percentile regions of CDTS, GERP and the overlap region of CDTS and GERP. CDTS, context-dependent tolerance score. GERP, Genomic Evolutionary Rate Profiling.

**Supplementary Figure S4.**
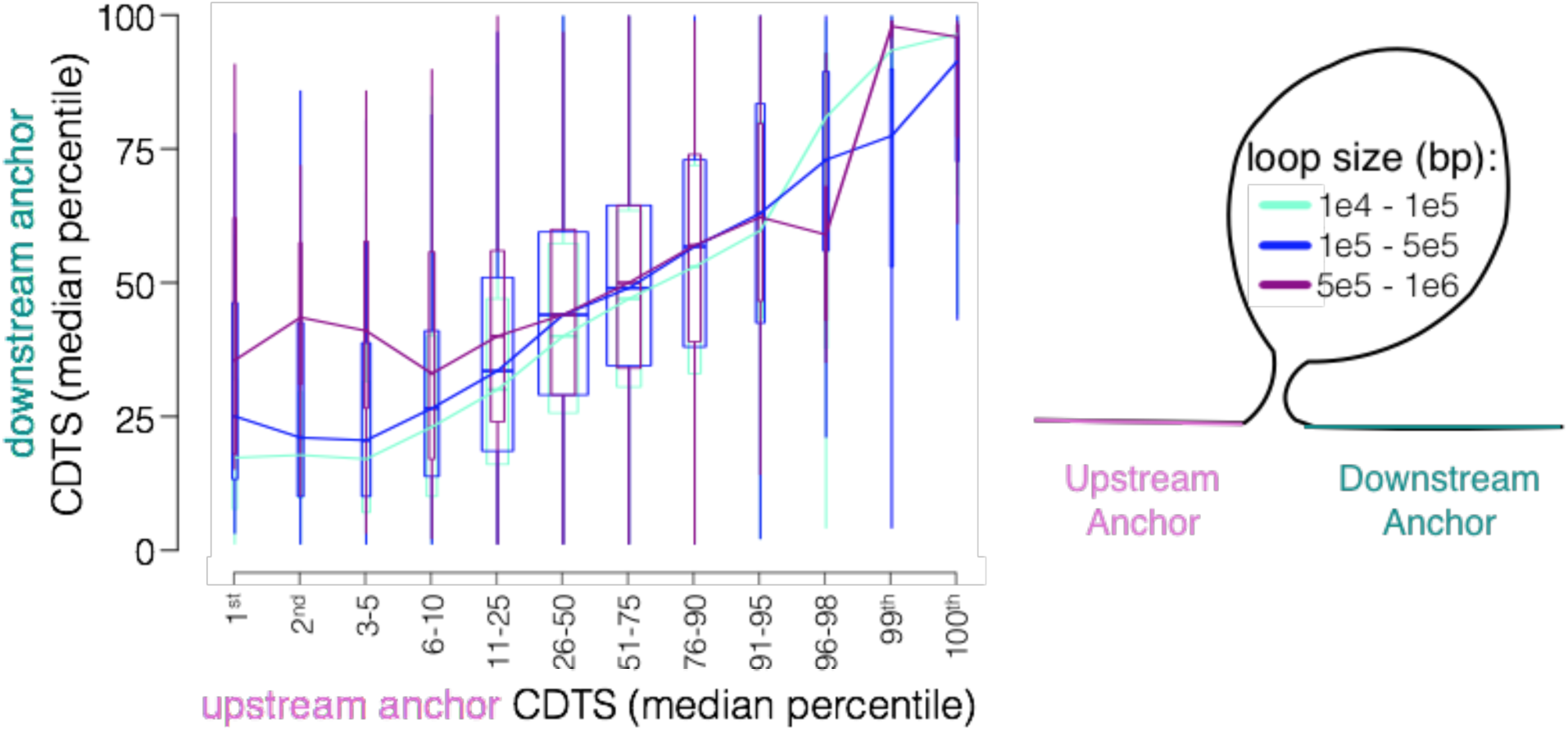
Shared conservation of distal anchor regions. Distal coordination of anchor regions grouped by the size of the loop. A chromatin loop is depicted in the right panel. The median CDTS is extracted for each anchor region and binned in percentile slices. The X- and Y-axes indicate the median CDTS values for the upstream and downstream anchor regions, respectively. The anchor regions surrounding a loop share CDTS values. The width of the boxplots is proportional to the number of anchors present in the respective CDTS percentile slice and with the respective loop size. The whiskers extend from the 10^th^ to the 90^th^ percentiles of the data. The box spans the interquartile range. Outliers are not displayed. The lines pass through the median of the respective boxplots.

**Supplementary Figure S5.**
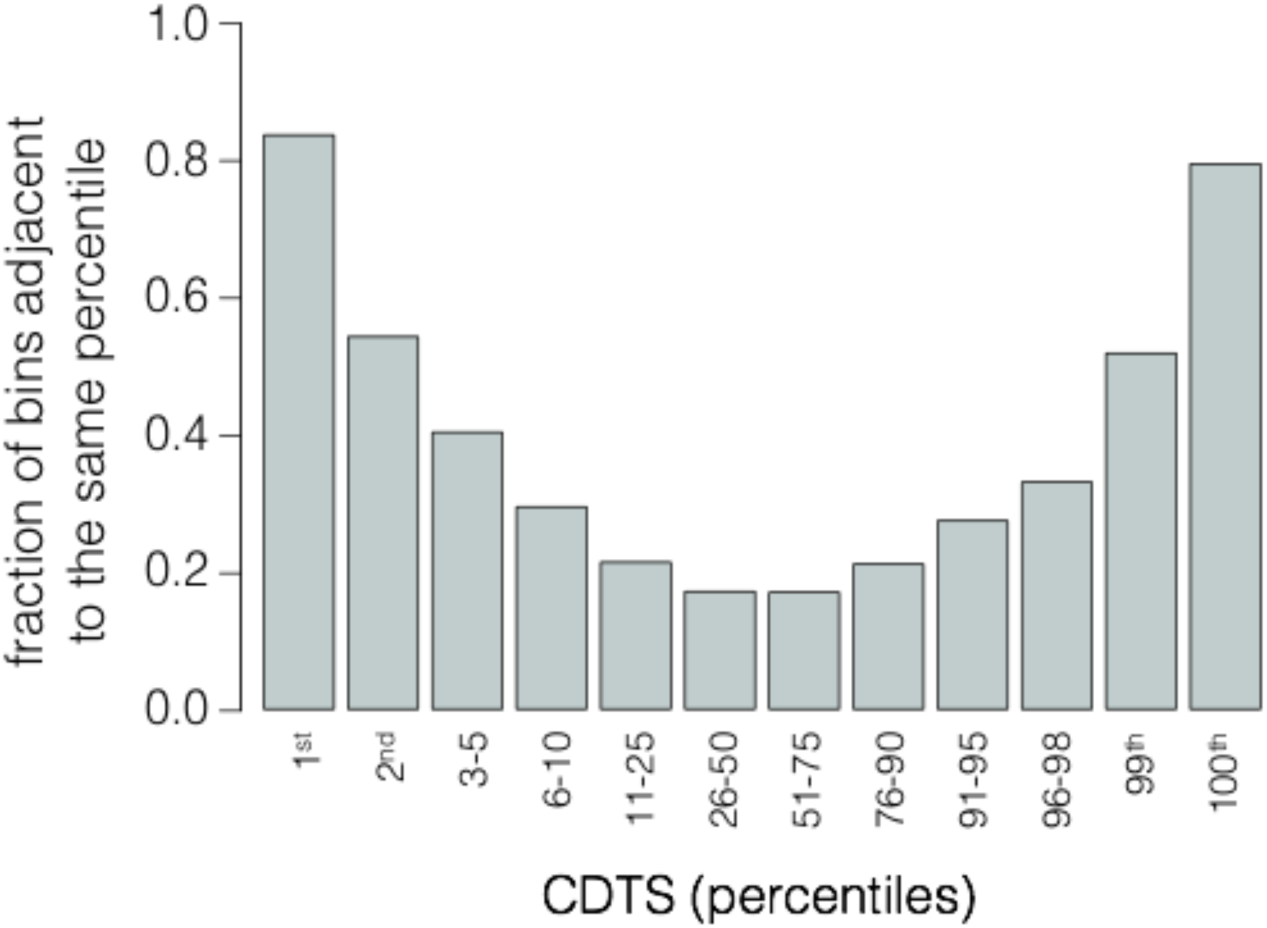
The barplot shows the fraction of bins that have an adjacent genomic bin in the same CDTS percentile. Bins in the 1st and 100st CDTS percentile tend to cluster together. CDTS, context-dependent tolerance score.

**Supplementary Figure S6.**
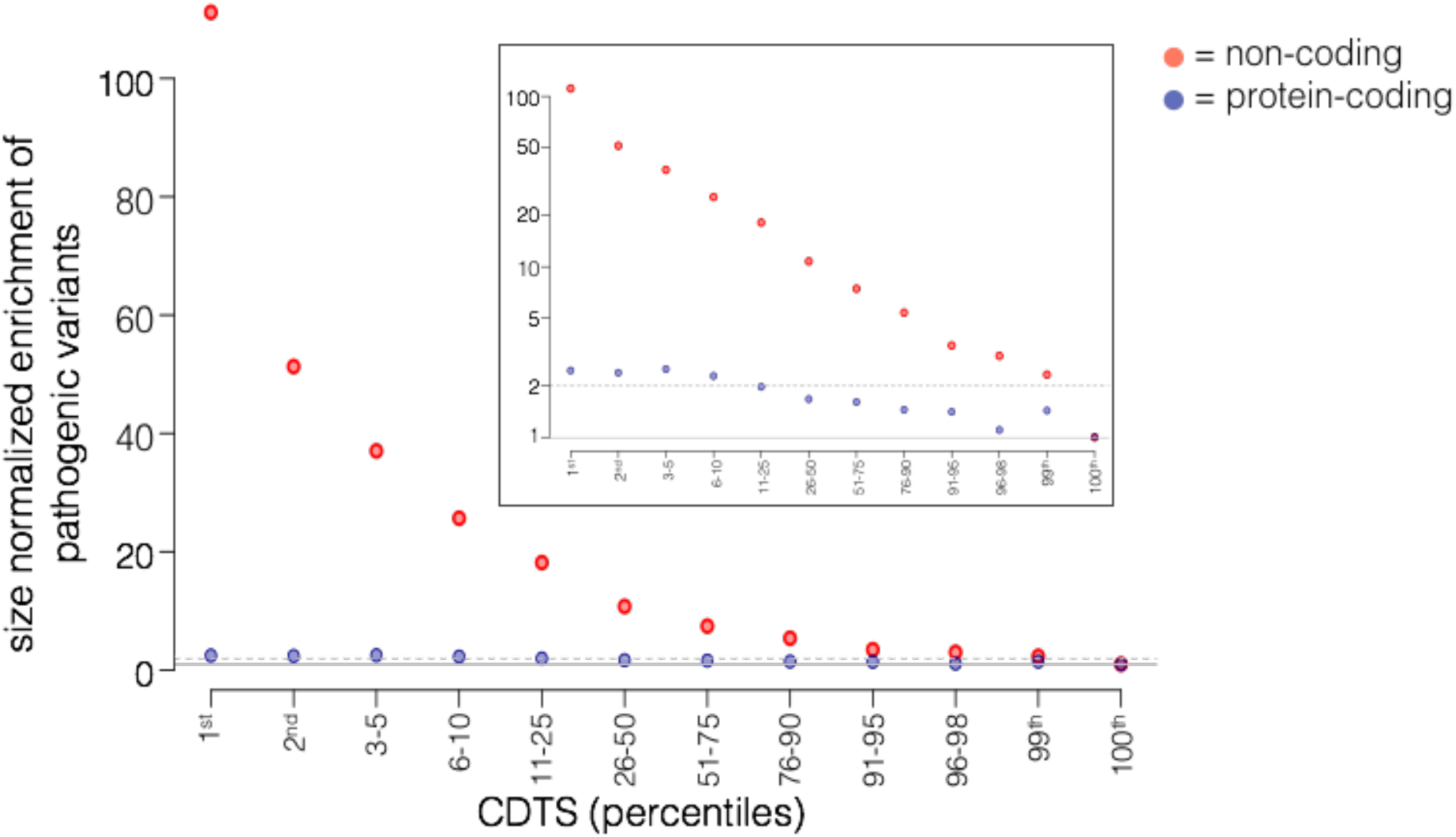
The distribution of pathogenic variants normalized by the size of the genomic element. The distribution of pathogenic variants across the different percentile slices is normalized by the size of protein-coding and non-coding regions in the respective percentiles slices. The relative enrichment is calculated with regards to the 100^th^ percentile. Protein-coding pathogenic variants are shown in dark blue; non-coding pathogenic variants in red. The total number of pathogenic variants are N=120,759 protein coding and N=14,092 non-coding variants. Exonic non-coding (e.g. lincRNA) are not displayed as only a very limited number of pathogenic variants were annotated (N=557). The inset figure is on a logarithmic scale. CDTS, context-dependent tolerance score.

**Supplementary Figure S7.**
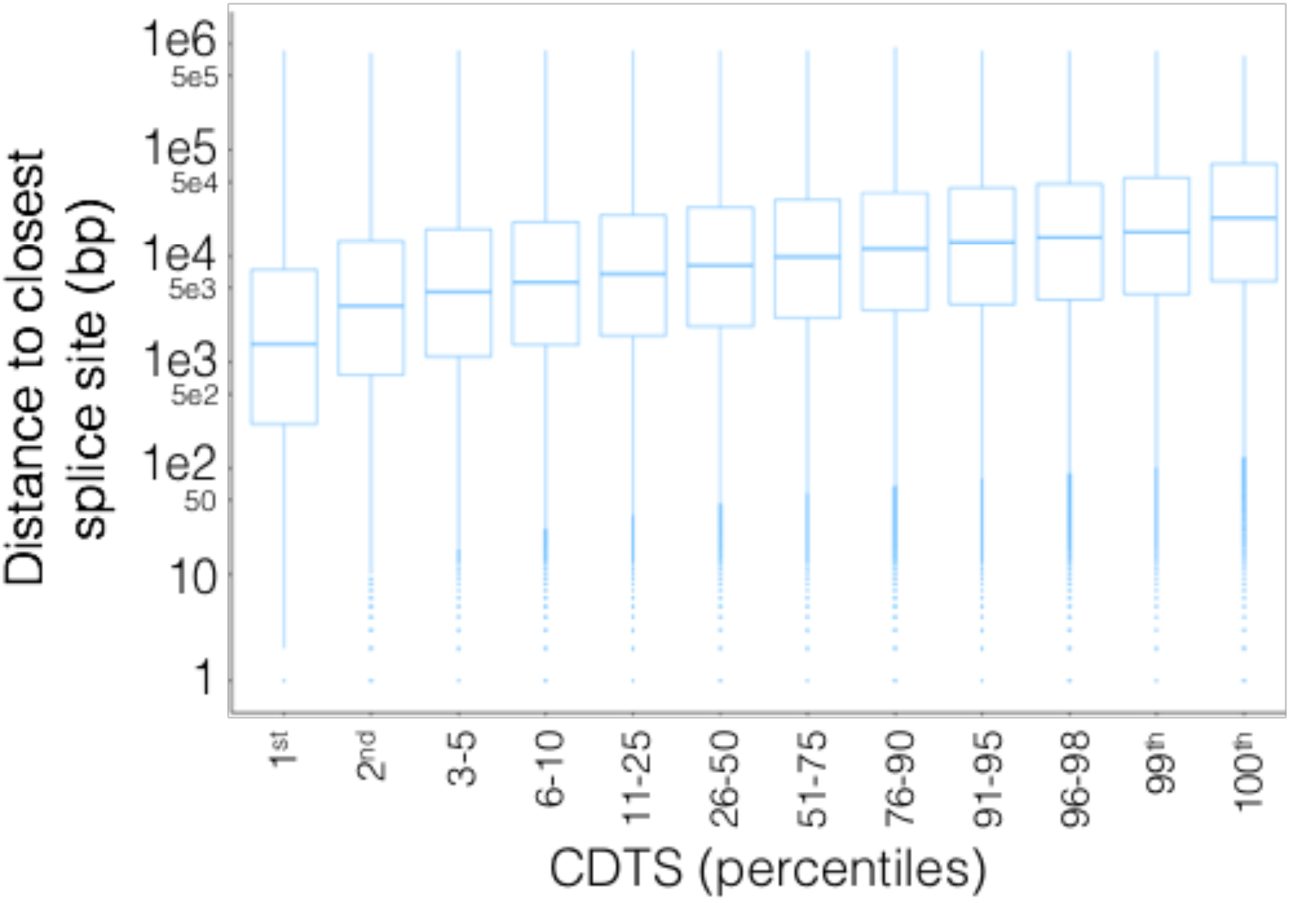
The boxplot displays for each intronic bin the distance to the nearest splice site. Intronic bins that have lower CDTS, appear to be closer to splice sites in general. The whiskers extend from the 10^th^ to the 90^th^ percentiles of the data. The box spans the interquartile range. The Y-axis is shown on a logarithmic scale.

